# Supracellular contractility in *Xenopus laevis* embryonic epithelia regulated by extracellular nucleotides and the purinergic G-protein coupled receptor P2Y2

**DOI:** 10.1101/2025.01.21.634176

**Authors:** Sagar D. Joshi, Timothy R. Jackson, Lin Zhang, Carsen Stuckenholz, Lance A. Davidson

## Abstract

Extracellular signals regulate epithelial homeostasis, cell fate and pattern cell behaviors during embryogenesis, wound healing, regeneration, and disease progression. Previous studies in our group found cell lysate from intentionally wounded embryos triggers a strong but transient contraction in neighboring epithelia, whether contiguous to the wound site or in non-wounded embryos. We previously identified extracellular ATP (eATP) as a possible candidate. Here we test additional candidates and find several nucleotides including ADP, UTP, and UDP also trigger contractility. Through a temporal and spatial screen of lysate activity, an inhibitor screen, and morpholino knock-down of candidate receptors, we find contractility is mediated by a G-protein coupled purinergic receptor P2Y2 (P2RY2). Activated P2RY2 triggers F-actin assembly and myosin II contractility. Knockdown of P2RY2 or overexpression of mutant G-protein effectors abrogate epithelial contractility when epithelia are exposed to eATP or lysate. We demonstrate that the major contributors to epithelial contractility in lysate are extracellular nucleotide triphosphates ATP and UTP, which are sensed by P2RY2 and transduced through G-proteins to contract the epithelium.

**Summary Statement:** Contractility of the *Xenopus* embryonic epithelium can be driven by extracellular nucleotides ATP or UTP and actuated by the G-protein coupled purinergic receptor P2Y2 (P2RY2).

**Highlights:**

- Extracellular nucleotides ATP and UTP can trigger epithelial contractility.
- Epithelia contract in response to (ATP ∼ UTP) > (ADP ∼ UDP) > ADO (adenosine).
- The purinergic G-protein coupled receptor P2Y2 is responsible for this contractile response by indirectly modulating actomyosin contractility.

## Introduction

While there is widespread belief that cell and tissue mechanical processes operating during vertebrate development are regulated at the supra-cellular scale, precise molecular pathways that function at that scale have been difficult to elucidate. To identify such factors, we aimed to investigate molecular pathways mediating tissue responses to wounding, particularly how responses are distributed across a field of cells that have not been directly injured (Joshi et al., 2010). These responses offered us the opportunity to identify signaling pathways that mediate this supracellular contractility. In this paper we describe a pathway involving extracellular ATP and a G-protein coupled receptor, P2RY2, that controls contractility in the embryonic epithelial of the frog *Xenopus laevis*.

## Results

### Cell lysate drives acute contraction of Xenopus embryonic epithelia

Acutely applied cell lysate can drive short term contraction of embryonic epithelia. We first confirmed results from previous study that lysate applied via perfusion can drive a transient contraction in *Xenopus* embryonic epithelia that lasts approximately 2 minutes (Fig. 1A; (Joshi et al., 2010)). Contraction begins 10 to 20 seconds after lysate is applied and continues for 30 to 90 seconds. Repeated perfusion of the same embryo at 30 second intervals demonstrates that the contraction response saturates (Fig. 1B; Supplemental Video 1); whereas pulsed delivery of lysate at 600 second intervals reveals that the response does not desensitize (Fig. 1C; Supplemental Video 2).

**Figure 1.**
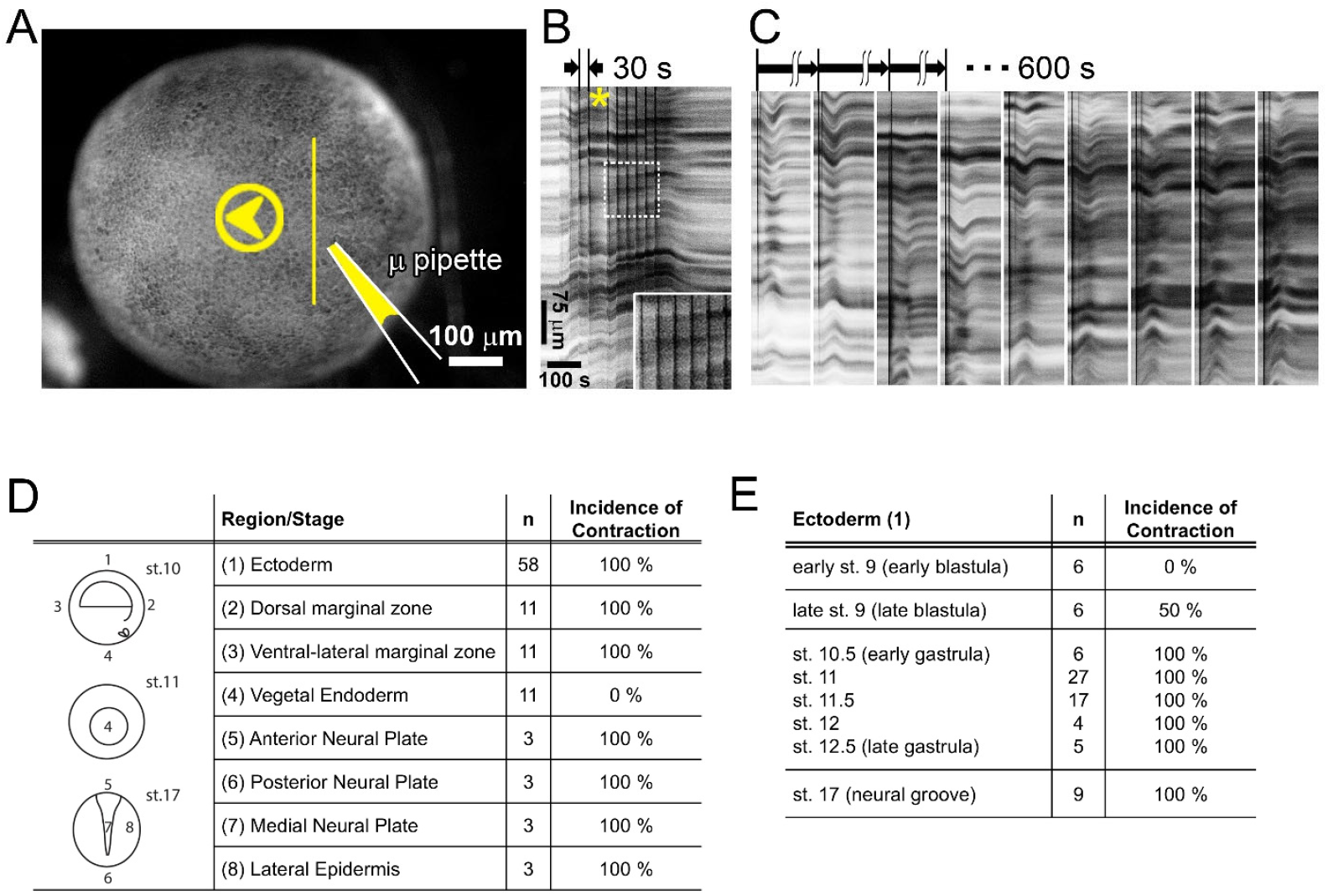
Spatiotemporal responses to lysate suggest lysate sensing does not desensitize and is developmentally regulated. A) A single frame from a timelapse demonstrating multiple pulses of lysate to the ectodermal surface of a gastrula stage embryo. Arrow head (yellow) shows the direction of lysate flow from the micropipette. Inset (dashed box) shows little change after multiple pulses. B) Kymograph from the timelapse sequence of 10 contractions in the same embryo (with a single 60 sec gap, asterisk; Supplementary Video 1). The surface is perfused with pulses of 60 nl lysate every 30 seconds. The embryonic epithelium recoils after the pulse train ends. C) Another representative set of kymographs shows the response of a single embryo perfused 9 times with 60 nl lysate every 600 seconds. The ectoderm contracts and relaxes after each perfusion (Supplemental Video 2; 9 sequences are concatenated). D) Contractile responses to perfused lysate are observed in all epithelial tissues in gastrula and neurula stages, except vegetal endoderm. E) Contractile responses in ectoderm perfused with lysate begin in late blastula stages and continue through neurulation.

### Contraction response is developmentally regulated

We next mapped temporal and spatial responses of developing embryos to lysate. We perfused different regions of developing embryos from late blastula (stage 9) to neural plate stages (stage 17). Animal cap ectoderm, the marginal zone, and all tissues near or within the neural plate epithelium respond, however, vegetal endoderm does not, even as nearby marginal zone cells are observed to contract (Fig. 1D). Next, we perfused lysate onto animal cap ectoderm in whole embryos ranging from early blastula to early neurula stages and found early blastula stage epithelia would not contract but that animal cap ectoderm and epidermis at later stages were fully responsive (Fig. 1E).The limited stage and tissue-specific responses suggest the molecular pathway mediating the contractile response is developmentally regulated.

### Lysate activates actomyosin contractility

Actomyosin contractility within the medio-apical cortex and apical junctions play a key role in epithelial homeostasis and morphogenesis (Sawyer et al., 2009; Martin et al., 2010; Agarwal and Zaidel-Bar, 2018; Blanchard et al., 2018; Miao and Blankenship, 2020). We previously found both the apical and basal cell cortex transiently assemble F-actin during exposure to lysate. To test whether F-actin and non-muscle myosin II mediate the epithelial contractile response to lysate, we used a panel of small molecules to inhibit actin polymerization (0.6 µM Latrunculin B; LatB; (Benink and Bement, 2005; Lee and Harland, 2007), to inhibit Rho Kinase (50 µM Y-27632; (Maekawa et al., 1999; Narumiya et al., 2000)) and to inhibit non-muscle myosin II heavy chain (100 µM Blebbistatin; BBS; (Straight et al., 2003; Lee and Harland, 2007)). We incubated early gastrula stage embryos for 30 to 60 minutes in each inhibitor, and perfused cell lysate over the animal ectoderm (Fig. 2A; Supplemental Video 3). Kymographs collected transverse to the flow stream at center of the contractile region (Fig. 2A’) can reveal both qualitative and quantitative changes in contraction magnitude and kinematics of the contraction event (Figs. 2B; (Joshi et al., 2010; Kim et al., 2014)). Notably, contractility at the same rate after F-actin polymerization or Rho Kinase activity inhibition but is frequently eliminated when myosin II is inhibited with BBS (Fig. 2C). In concordance with the role of actomyosin in cell contractility, all treatments significantly reduced the strength, or absolute magnitude of the contractile response (Fig. 2D). BBS produced the largest reduction in the magnitude of the contraction (even abolishing the response in one clutch) and increased the delay between stimulation and onset of the contractile response in cases where a contraction was observed (Fig. 2E). While BBS did not significantly reduce the time to peak (Fig. 2F), the inhibition of myosin II contractility shortened the duration of contractions (Fig. 2G). Inhibition of F-actin polymerization and Rho Kinase both significantly reduced contraction strength significantly (Fig. 2D), but were less effective than direct myosin II inhibition. Remarkably, disruption of F-actin by LatB to the point where epithelial integrity is lost and cell-cell junctions begin to rupture (see asterisks in Fig. 2A) only moderately reduced contractility. Thus, actomyosin contractility is a key target of the cellular response to acute lysate stimulation.

**Figure 2.**
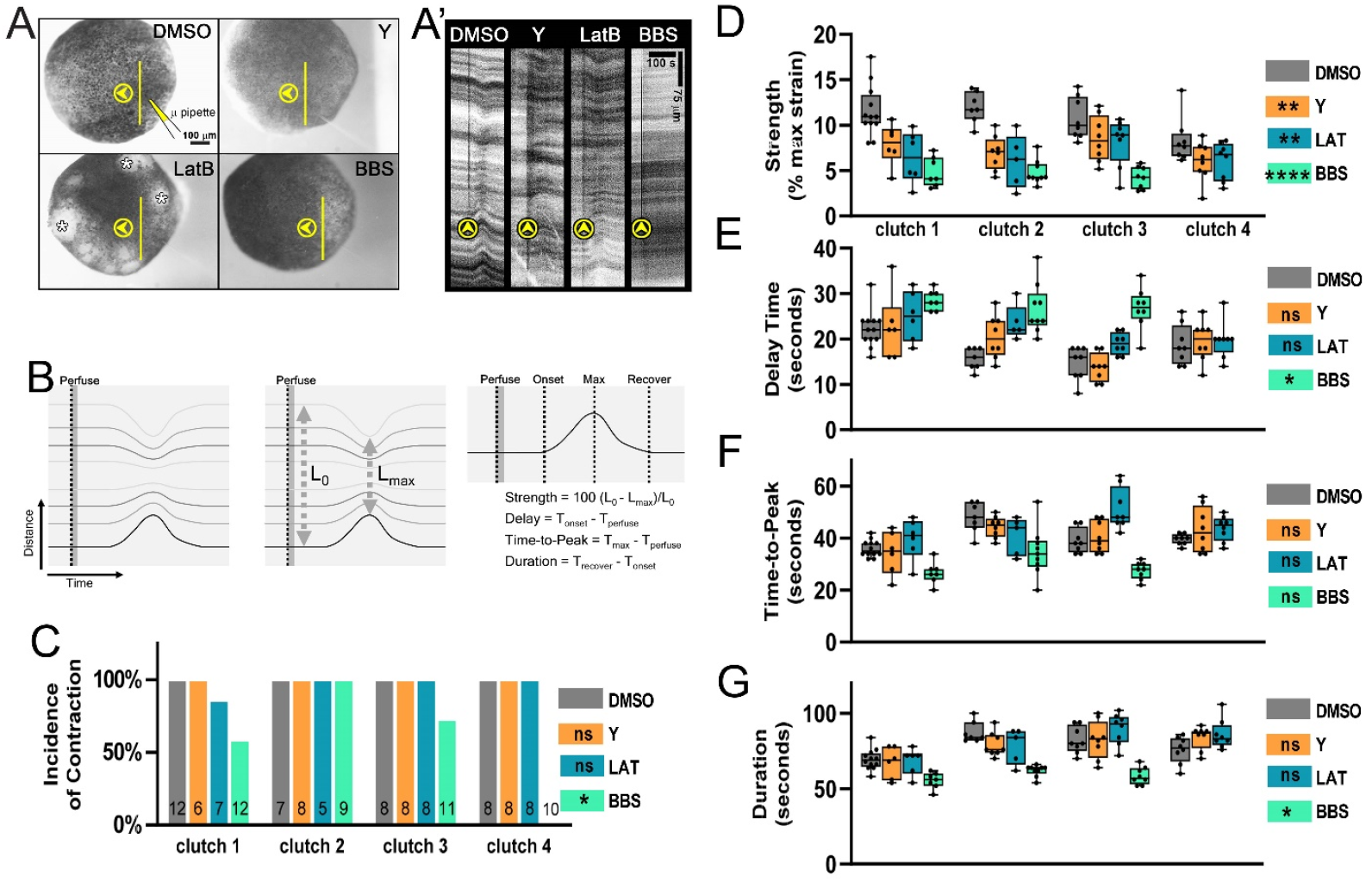
F-actin and myosin II are involved in contractile response to ATP. A) Single timelapse frames of embryos incubated in DMSO (carrier control), Y-27632 (Y), latruculin B (LatB), or blebbestatin (BBS) prior to being exposed to a single 60 nl pulse of lysate. Arrow heads indicate direction of perfusion flow (see Supplemental Video 3). A’) Kymographs collected from timelapses along lines (in A) show contractions for each condition. Vertical arrowheads indicates time of perfusion. Multiple lesions in the epithelium appear after treatment with LatB (asterisks). B) Schematic kymograph showing a mock contraction with lines indicating pigment features displacing over time and their analysis under each condition: C) Incidence of contraction (percent), D) Strength, E) Delay Time, F) Time-to-Peak, and G) Duration of the contraction. Note: Cases where contractions are absent are not included in (D) through (G). Statistics for (C) through (G) indicate significance from DMSO control condition (ns, not significant; *, p< 0.05; **, p< 0.01; ****, p < 0.0001).

### Extracellular nucleotides ATP, ADP, UTP and UDP, present in lysate, induce contractions

We tested several candidates in order to identify what factor or a group of factors induced actomyosin contractility. Using a perfusion-based candidate screen, we confirmed extracellular ATP (eATP) as a contractility agonist (Kim et al., 2014) and that UTP was also capable of inducing contractions (Table 1). Extracellular ADP, adenosine (ADO) and UDP also induce contractility but only at concentrations of 40 µM or higher. Additionally, the energy source of eATP is not required to drive contractions since ATPγS, a non-hydrolysable analog of ATP can also drive contractility. Furthermore, treatment of ATP with shrimp alkaline phosphatase (SAP), a nucleotide hydrolyzing enzyme, abrogated the ability of 40 µM ATP to induce contractions but did not reduce contractions driven by ATPγS. To test whether eATP, or other nucleotides in lysate were triggering contractility, we incubated lysate with SAP and found it could not induce contractions (Supplementary Table 1), similar to embryo culture media, salts, sodium glutamate, and acetylcholine. Thus eATP and a limited set of other nucleotides present in lysate are sufficient to induce contractions with relative selectivity: ATP≈TP > ADP≈DP > ADO. Since ATP is by far the most abundant nucleotide in *Xenopus* egg cytoplasm, approximately 1 mM range (Woodland and Pestell, 1972), we focused this study on the effects of extracellular ATP.

**Table 1.** Testing nucleotides and nucleosides for ability to induce epithelial contraction.

| Candidate Factors |  |  | # of embryos | %<sup>1</sup> | Contraction Magnitude<sup>2</sup> |
| --- | --- | --- | --- | --- | --- |
| **Embryos perfused with Nucleotides/ Nucleosides** | Adenosine triphosphate (ATP) | 400 µM | 6 | 100 | ++ |
|  |  | 40 µM | 6 | 100 | ++ |
|  |  | 0.4 µM | 18 | 72 | + |
|  | Adenosine diphosphate (ADP) | 40 µM | 4 | 100 | + |
|  | Adenosine (ADO) | 400 µM | 8 | 75 | + |
|  |  | 40 µM | 5 | 0 | - |
|  | Cyclic adenosine monophosphate (cAMP) | 40 µM | 5 | 0 | - |
|  | Cytidine triphosphate (CTP) | 40 µM | 8 | 0 | - |
|  | Guanosine triphosphate (GTP) | 40 µM | 6 | 0 | - |
|  | Uridine triphosphate (UTP) | 40 µM | 6 | 100 | ++ |
|  | Uridine diphosphate (UDP) | 40 µM | 6 | 100 | + |
|  |  | 0.4 µM | 5 | 0 | - |
|  | Thymidine triphosphate (TTP) | 40 µM | 8 | 0 | - |
|  | ATPγS | 2 mM | 14 | 100 | + |
|  |  | 1 mM | 7 | 100 | + |
|  |  | 100 µM | 13 | 15 | + |
|  | SAP treatment | ATP (40 µM) | 6 | 0 | - |
|  |  | ATPγS (1mM) | 10 | 100 | + |

<sup>1</sup>Incidence of contraction may be any magnitude (++ or +) in response to perfusion. <sup>2</sup>Contraction magnitude is qualitatively assessed comparing nucleoside/nucleotide perfusions of embryos with an observable contraction. Contraction magnitude is categorized with respect to 40µM ATP or Lysate: (++) contraction is equal or stronger, (+) contraction is weaker, (-) contraction not observed.

### The purinergic receptors mediate effects of eATP on contractility

Extracellular ATP is sensed by cells via two families of purinergic cell surface receptors, G-protein coupled receptors (GPCRs) of the P2Y-family (P2YR) and L-type Ca^++^ channel coupled receptors of the P2X-family (P2XR) (Abbracchio and Burnstock, 1994; Giuliani et al., 2019). Both receptor families elevate intracellular Ca^++^ in response to eATP. In *Xenopus*, eATP can trigger Ca^++^ influx in embryonic ectoderm (Kim et al., 2014) as well as driving bursts of F-actin assembly in the apical cell cortex (Arnold et al., 2019). To test the potential role of P2XR coupled Ca^++^ channels, we incubated embryos in the L-type calcium channel antagonist Lanthanum chloride (4 mM)(Lettvin et al., 1964), perfused lysate, and found ectoderm contracted with the same frequency and strength as controls (Supplementary Table 2). To further discriminate between the potential roles of P2Y- and P2X-families of eATP receptors, we perfused lysate onto embryos incubated in a competitive inhibitor of P2XR, pyridoxal-phosphate-6-azophenyl-2’,4’-disulfonate (PPADS, 100 µM; Supplementary Table 2) (Lambrecht et al., 1992; Ziganshin et al., 1993). When perfused with high concentrations of eATP (400 µM) ectoderm continued to contract, however, at lower eATP concentrations (0.4 µM), PPADS eliminated contractions. Since inhibition of P2X receptors failed to block eATP induced contractility we focused our candidate screen on the P2Y-family of receptors.

### Purinergic receptor P2Y2 mediates contractility response to extracellular ATP

The *Xenopus laevis* genome (Session et al., 2016) encodes 11 members of the P2Y-family. To narrow the set of candidates, we used RT-PCR to screen expression of P2Y receptors that shared ligand specificity (ATP, UTP, ADP, UDP, or ADO) (Supplementary Table 3) to determine which receptors are expressed at the stages or in tissues with visible contractile responses (Fig. 1C and D). Additionally, we ruled out roles for two P2Y-family members, P2Y8 and P2Y11, that had been previously described. One family member, P2Y8, appears associated with *Xenopus* neural patterning (Bogdanov et al., 1997), but is not expressed in the late blastula or early gastrula. Another family member, P2Y11 has been associated with sporadic contraction (Shindo et al., 2010) and cAMP signaling (Devader et al., 2007) during *Xenopus* gastrulation, but this receptor was not expressed in animal ectoderm. Of the remaining candidates, we found P2RY2 isoalleles P2RY2.L and P2RY2.S are expressed in the early embryo with a temporal pattern that closely aligns with gastrulation (Session et al., 2016). We further confirmed the expression of P2RY2 by RNA in situ hybridization and found that it was absent in early- and mid-gastrula endoderm but expressed in ectoderm (Fig. 3A), consistent with the spatial pattern observed in perfusion studies (Fig. 1D).

**Figure 3.**
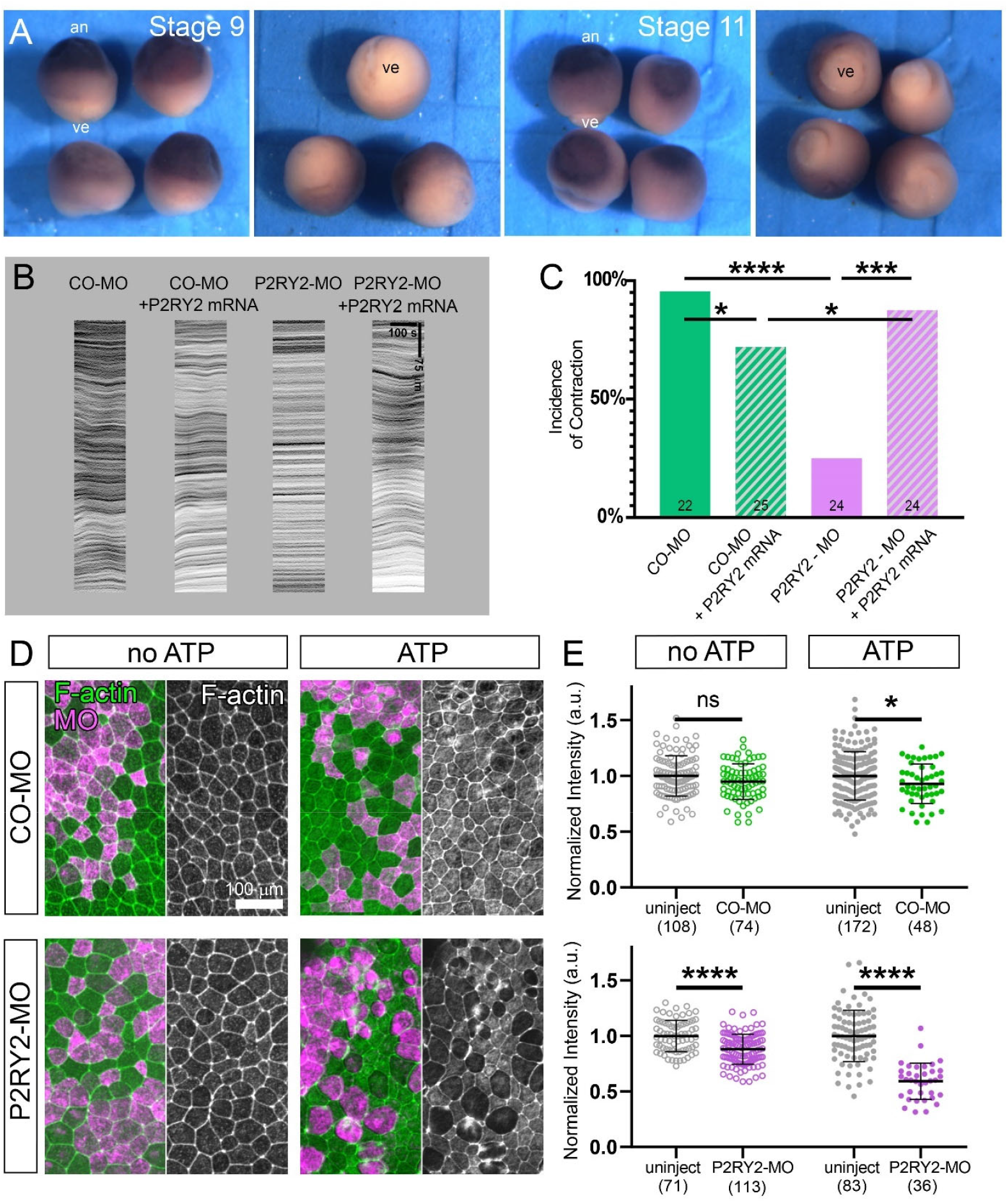
Gene expression and knock-down of P2Y2 Receptor. A) RNA in situ hybridization gene expression of P2RY2 at late blastula (stage 10) and mid-gastrula (stages 11). Highest expression is seen in the animal pole (an) with the lowest in the vegetal endoderm (ve). (B) Representative kymographs of contractility of morpholino controls, knock-downs, controls, and rescues after perfusion with 4nL of 400 µM ATP. For knockdown of P2RY2, embryos were injected with either 32 ng of a control morpholino oligomer (CO-MO) or 16 ng each of two morpholino oligomers targeting translation initiation sites of the two alloalleles, P2RY2.L and P2RY2.S (P2Y2-MO). Rescue was tested in embryos co-injected with 2 ng of a morpholino-resistant P2RY2.L mRNA. C) The incidence of contractility for knockdown and rescue. D) P2RY2 knockdown effects on actomyosin accumulation after exposure to eATP. The left side of each panel shows F-actin stained (green) and a scattered population morpholino expressing cells (rhodamine dextran; magenta) alongside a grayscale image of the F-actin channel. E) Medioapical F-actin intensity was measured in knockdown and control morpholino injected explants and compared between uninjected and injected, e.g. morpholino injected, cells. The intensity of each ROI, in both uninjected and MO-injected cells, was normalized to the mean F-actin intensity of the uninjected cells in the same explant. Results for each condition are pooled from 3 to 4 explants and cell numbers are listed below each data column. Statistics for (C) and (E) indicate significance on the Incidence of Contraction and the Normalized Intension of F-actin, respectively (ns, not significant; *, p < 0.05; ***, p> 0.001; ****, p>0.0001).

We tested the necessity of P2RY2 in eATP-induced contractility by simultaneous knockdown of both P2RY2 alloalleles. We co-injected two morpholino oligonucleotides designed to block the translation initiation sites of both P2RY2.L and P2RY2.S (P2RY2-MO) into animal blastomeres at the 8-cell stage (32 ng total/embryo). We found P2RY2-MO strongly inhibited contraction responses to eATP perfusion, with only 25% of P2RY2-MO injected embryos exhibiting a contractile response compared to 93% of control morpholino (COMO) injected embryos (Fig. 3B & C). Furthermore, co-injection of a morpholino-resistant mRNA encoding P2RY2.L successfully rescued eATP-induced contractility in P2RY2-MO injected embryos with 88% of embryos responding to eATP.

### Purinergic receptor P2Y2 regulates apical F-actin levels, increasing levels after eATP

Perfused eATP and lysate can drive rapid remodeling of the actin cytoskeleton, enhancing apical (Arnold et al., 2019) and transiently depolymerizing and remodeling basal networks (Joshi et al., 2010). To test whether apical F-actin assembly is regulated by P2RY2, we injected COMO or P2RY2-MO contralaterally together with a Rhodamine-dextran lineage label into the animal pole at 4- or 8-cell stages. We raised the injected embryos to early gastrula stages, explanted animal caps onto fibronectin coated substrates (Joshi and Davidson, 2010), and added ATP or culture media to the samples. After 30 minutes, explants were fixed and stained with phallacidin to quantify F-actin intensity in both injected and uninjected cells. Due to sample-to-sample variation in fixation and staining, we normalized levels of F-actin to levels in neighboring uninjected cells (Fig. 3D & E). In the absence of ATP, ectoderm cells expressing P2RY2 -MO showed 11% reduction in F-actin compared to uninjected cells in the same explant. The impact of the P2RY2 knock-down was pronounced after P2RY2-MO expressing cells were exposed to eATP, resulting in a 40% reduction in F-actin intensity in P2RY2-MO expressing compared to uninjected cells. The COMO injected cells also saw a 10% reduction in F-actin compared to uninjected cells. Together with the ability of actomyosin inhibitors to modulate contractile responses to lysate, our knock-down findings support the central role of P2RY2 in eATP-induced contractility via F-actin remodeling.

Once activated by eATP, P2RY2 may drive contractility through direct association with the cytoskeleton (Yu et al., 2008; Orchard et al., 2014; Haenig et al., 2020) or via G-protein activation of downstream actomyosin effectors such as RhoA or Phospholipase C (Erb and Weisman, 2012; Flock et al., 2017). To test the role of G-protein signal transduction pathways, we co-injected mRNA encoding prenylation-deficient forms of Gγ-3 and Gγ-5 in which the prenylation sequence CAAX has been mutated to SAAX (Mulligan et al., 2010). SAAX forms of Gγ proteins bind Gβ proteins, sequestering them in the cytoplasm, and inhibit Gβγ localization and activation by GPCRs at the plasma membrane. Animal cap epithelia expressing both Gγ-3-and Gγ-5-SAAX proteins from injected mRNAs exhibited 40 to 60% of the contractility strength of controls (Supplementary Fig. 3A & B). The reduction of contractility by Gγ-SAAX over-expression demonstrates involvement of a G-protein mediated pathway.

Extracellular nucleotides, including ATP and UTP released from wounded epithelia can regulate actomyosin contractility. Here, we demonstrate diffusible extracellular ligand ATP (eATP) drives contractility in embryonic epithelium through P2RY2. eATP and its cellular receptor P2RY2 are well known regulators of embryonic and adult tissue mechanics (Burnstock and Verkhratsky, 2012; Verkhratsky and Burnstock, 2014; Mikolajewicz et al., 2019). eATP and P2RY2 are involved in mammalian deciduation and implantation (Gu et al., 2020). eATP release after mechanical stimulation is well characterized in the cardiovascular system where it plays a role in flow sensing by endothelial (Milner et al., 1990) and red blood cells (Wan et al., 2011), triggering valvulogenesis (Fukui et al., 2021), and is involved in the cellular stress response (Galluzzi et al., 2018). eATP and P2RY2 additionally regulate tissue responses to mechanical stimulation of lung epithelia (Homolya et al., 2000), cancer (Burnstock and Di Virgilio, 2013), and the central nervous system (Burnstock, 2008). The activity of eATP and P2RY2 suggest a role in *Xenopus* epithelial homeostasis and morphogenesis. The results demonstrate actomyosin contractility, likely driven by eATP could be a key mechanism for controlling the contractile forces that shape tissues and drive organogenesis. This suggests a linkage between biochemical regulation of actomyosin and supracellular control of tissue-scale movements.

## Methods

### Embryo and explant culture, microsurgery, histology, morpholinos, and RNA *in situ* hybridization

Eggs were obtained from female *Xenopus laevis* frogs, fertilized, dejellied in 2% Cysteine solution (pH 8.4), and cultured in 1/3× Modified Barth’s solution (MBS) following standard methods (Kay and Peng, 1991) in accordance with the animal use protocols of the University of Pittsburgh. Embryos were staged appropriately (Nieuwkoop and Faber, 1967) and vitelline membranes were removed using forceps. Animal cap explants were microsurgically isolated and cultured in Danilchik’s For Amy solution (DFA; (Sater et al., 1993)) and cultured on fibronectin-coated glass substrates (Joshi, 2011). Explants processed for F-actin localization were fixed with 4% paraformaldehyde and 0.25% glutaraldehyde in PBST (1x PBS with 0.1% Triton X-100) for 15 minutes at room temperature. After washing, the samples were incubated with BODIPY-FL–phallacidin (1:800) to visualize F-actin. Morpholino oligonucleotides were designed with GeneTools Inc. (Philomath, OR) to block the translational start sites of P2RY2.L: TTCTGGGTCTTCAAACACATTCATC; P2RY2.S: CTTGGTCTCCAGACAAATTCATTTT.

We note that P2YR2 genes consist of a single exon, precluding design of splice acceptor morpholinos. Embryos were processed for gene expression analysis using standard RNA in situ hybridization protocols (Harland, 1991). Probe cDNA were amplified from stage 10 libraries with the following primers:

P2RY2.L Forward primer: GGTTGAGACTACAAAACCAGAA,

P2RY2.L Reverse primer: CTATGTTACCTCATGGAGGTTG;

P2RY2.S Forward primer: GGTTAAAACAACAAAACGAGAAAG,

P2RY2.S Reverse primer: GGTATATGTAGATTTACCTACAGCC.

### Lysate, ATP, and Compound Perfusion

The perfusion and lysate preparation methods have been described previously (Joshi et al., 2010). Lysate and dilute perfusion compounds were freshly prepared on the day of the experiment. Salts, bioactive compounds, and nucleotides/ nucleosides were diluted in 1/3× MBS (See Table 1 and Supplementary Table 2 for tested compounds). All compounds were stored and utilized according to the product guidelines (Sigma-Aldrich, St. Louis MO). To visualize cell lysate and other perfused solutions, 3 μl black non-waterproof ink (Higgins Fountain Pen India Ink; Utrecht Art, Cranbury NJ) was added to each 100 μl stock perfusate. Since contractile responses are variable in embryos from one mating to the next (e.g. “clutch”), for instance the strength of contraction varies from 9 to 12%, we compare results from multiple clutches.

### Image acquisition, analysis, and statistical analysis

Still images, time-lapse sequences of developing embryos, and time-lapse sequences of perfusion experiments were recorded as described previously (Joshi and Davidson, 2010) using a CCD camera (Scion Corp., Frederick, MD) mounted on a dissecting stereomicroscope. Image acquisition and subsequent analysis of time-lapses sequences were carried out using custom-written macros and plug-ins for image analysis software (MicroManager 1.4 and 2.0, (Edelstein et al., 2014); ImageJ ver. 1.54, (Schneider et al., 2012). To generate kymographs, a line region-of-interest spanning the center of the embryo was selected and resliced over the full timelapse sequence. Contrast was enhanced and each kymograph was manually categorized for incidence into strong contraction, weak contraction, or no contraction. Measurements of contraction strength, delay time, time-to-peak, and duration were quantified as previous (Joshi et al., 2010). Tests for statistical significance of treatments were determined with simple and paired T-tests, 2-WAY ANOVAs, or Mann-Whitney U-tests (Sokal and Rohlf, 1987) using commercial statistical software (GraphPad Prism v. 10 and SPSS v. 25).

## Acknowledgements

We would like to thank members of the MechMorpho Lab for their support and lively discussion as this project evolved. Logic is the beginning of wisdom, not the end. This work was supported by grants from the NIH (R01 and R37 HD044750, R21 HD106629, and R56 HL134195) and the University of Pittsburgh Swanson School of Engineering. TRJ was additionally supported by the Cardiovascular Bio-engineering Training Program (NIH NHLBI T32 HL076124).

