## Supplemental Data for "Supracellular contractility in *Xenopus laevis* embryonic epithelia regulated by extracellular nucleotides and the purinergic G-protein coupled receptor P2Y2"

**Supplemental Tables:**

**Supplementary Table 1. Testing compounds for ability to induce epithelial contraction.**

| | Candidate Factors | # of embryos | %[1] | Contraction Magnitude[2] |
|---|---|---|---|---|
| **Controls** | Lysate (2 embryos lysed per 100 µl 1/3X MBS) | 49 | 100 | + + |
| | 1/3X MBS | 6 | 0 | - |
| | 20% Ink (Higgins Fountain Pen India Ink) in 1/3X MBS | 6 | 0 | - |
| **Salts** | Potassium chloride (KCl, 1M) | 6 | 0 | - |
| | Sodium chloride (NaCl, 1M) | 6 | 0 | - |
| | Calcium chloride ($CaCl_2$, 1M) | 6 | 0 | - |
| | Magnesium chloride ($MgCl_2$ , 1M) | 5 | 0 | - |
| **Bioactive Compounds** | Sodium glutamate ($C_5H_8NNaO_4$, 1M) | 3 | 0 | - |
| | Acetylcholine (3.1 mM) | 3 | 0 | - |
| **Treatments** | Lysate - heat treated (65 °C, 30 minutes) | 9 | 100 | ++ |
| | Lysate - shrimp alkaline phosphatase treated (SAP, 1U/ 10 µl) | 5 | 0 | - |

[1]Incidence of contraction may be any magnitude (++ or +) in response to perfusion. [2]Contraction magnitude is qualitatively assessed for trial perfusions of embryos with an observable contraction. Contraction magnitude is categorized with respect to 400 µM ATP or Lysate: (++) contraction is equal or stronger, (+) contraction is weaker, (-) contraction not observed.

**Supplementary Table 2. Testing inhibitors for ability to block epithelial contraction by lysate or ATP.**

| | Candidate Inhibitors | Perfused with | # of embryos | %[1] | Contraction Magnitude[2] |
|---|---|---|---|---|---|
| **Embryos incubated in compounds for 2 hours** | Lanthanum chloride (4 mM; L-type Calcium channels; (Lettvin et al., 1964)) | Lysate | 20 | 100 | ++ |
| | Pyridoxalphosphate-6-azophenyl-2',4'-disulphonic acid (PPADS 100 μM; P2X/P2Y; (Lambrecht et al., 1992)) | ATP (400 μM) | 6 | 100 | ++ |
| | | ATP (40 μM) | 8 | 100 | ++ |
| | | ATP (4 μM) | 4 | 100 | ++ |
| | | ATP (0.4 μM) | 8 | 0 | - |

[1]Incidence of contraction may be any magnitude (++ or +) in response to perfusion. [2]Contraction magnitude is qualitatively assessed for trial perfusions of embryos by comparison with an observable contraction. Contraction magnitude is categorized with respect to 400 μM ATP or Lysate: (++) contraction is equal or stronger, (+) contraction is weaker, (-) contraction not observed.

**Supplementary Table 3. PCR Expression Profile of P2YR.**

| | Primer 1 | Primer 2 | St 0 | St 8 | St 10 | St 10 ecto |
|---|---|---|---|---|---|---|
| P2Y1.L/S | CTCAAAGCACTTTGCTGGCAAGTG | GCTTGTGTCCCCATTCTGCTTG | - | - | - | - |
| P2Y2.L | CGTCCTCCTGCCGGTGTCGT | GCACTGGCCAGGGGTCTGGT | - | + | + | + |
| P2Y2.S | GACAACTGGCCCTTCGGCGT | CAAGGCAGCTGTTGGCGCTG | - | + | + | + |
| P2Y4.L | TGCTGCCGGTCTCCTACAGTGC | TTGATCCCAAGCAACCGGGCA | - | - | - | - |
| P2Y6.L | ACGTGTGCTCCCTCCCGCTC | ACCAGGCAGGGCACATCGTCC | + | + | + | + |
| P2Y6.S | CCCGCACGGCCATCTACACC | GACAGGGCCGGAGGGCTCA | + | + | + | + |
| P2Y8.L | TGGCCTGGGCAGCCTTCATC | GTTCACAGCTGCGATGGGTGC | - | - | - | - |
| P2Y8.S | AGGCACCTTTGCACGCTGGT | CCTGCCGCTTACAGCTGCGAT | - | - | - | - |
| P2Y10.L | CACATGGCCGTTCGGACGCT | AAGGCGCCCACGCATCACTG | - | - | - | + |
| P2Y10.S | ACCGCAAGTGGGCGTTTGGA | TCCGTGCTTCGCGACCTCAT | - | - | - | + |
| P2Y11.L | GGAGTTTGTGGTGTCTGTGCTCGG | ACAGGGGCAGAGATGGCAGC | - | + | + | + |
| P2Y11.S | TGGGCCTGGGGGCTCAAAGA | AGCAGCGGGTGGATACAGGGA | - | + | + | + |
| P2Y12.L | TGGGTTGGCAGTGCGTGTGT | GGCCGGCCTTTTTCTCCACA | + | + | + | + |
| P2Y12.S | TTCGAAACTCGGGCTATGGAAGC | TCCACATGGCCAGCAGCGAG | - | - | - | - |
| P2Y13.L | TGGCTCCTCATTGCTCCCCA | AGCAGCCAGCCAAAGCGTGA | - | - | - | - |

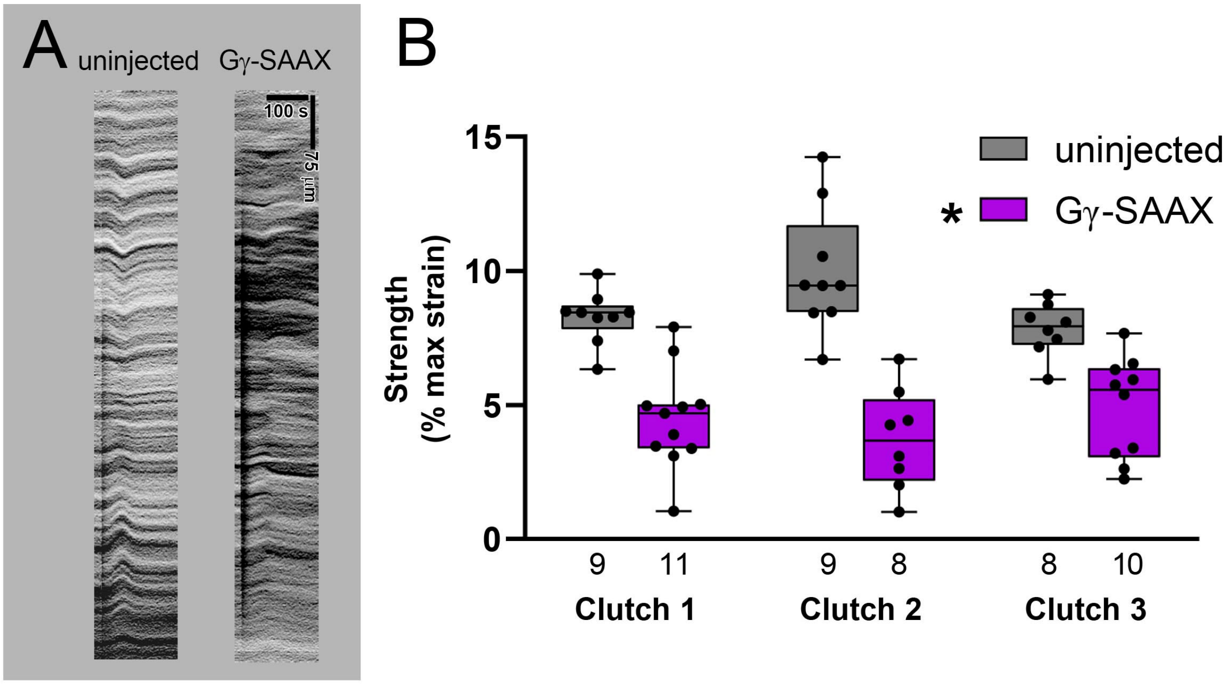


**Supplemental Figure S1. Gγ-SaaX proteins reduce contractility.** A) Kymographs of Xenopus ectoderm expressing Gγ-SaaX dominant negative constucts shows reduced contractility when perfused with 40 nl of 40 μM ATP. B) The strength of contractility is measured as in Fig. 2B. The number of embryos in each set is indicated below the data column. Statistics in (B) indicates significance of the Strength of Contraction (*, $p < 0.05$).
